# Enzymatic assembly of small synthetic genes with repetitive elements

**DOI:** 10.1101/2022.11.23.517630

**Authors:** Michael T. A. Nguyen, Martin Vincent Gobry, Néstor Sampedro Vallina, Georgios Pothoulakis, Ebbe Sloth Andersen

## Abstract

Gene synthesis efficiency has greatly improved in recent years but is limited when it comes to repetitive sequences and results in synthesis failure or delays by DNA synthesis vendors. This is representing a major obstacle for the development of synthetic biology, since repetitive elements are increasingly being used in design of genetic circuits and design of biomolecular nanostructures. Here, we describe a method for the assembly of small synthetic genes with repetitive elements: First, a gene of interest is split *in silico* into small synthons of up to 80 base pairs flanked by Golden Gate-compatible overhangs. Then, synthons are made by oligo extension and finally assembled into a synthetic gene by Golden Gate Assembly. We demonstrate the method by constructing eight challenging genes with repetitive elements e.g., multiple repeats of RNA aptamers and RNA origami scaffolds with multiple identical aptamers. The genes range in size from 133 to 456 base pairs and are assembled with fidelities of up to 87.5 %. The method was developed to facilitate our own specific research but may be of general use for constructing challenging and repetitive genes and thus a valuable addition to the molecular cloning toolbox.

## Introduction

Advances in DNA synthesis technologies have been pivotal for synthetic biology research. With the development of DNA assembly methods such as restriction-ligation^1^, polymerase chain assembly^2^, ligation-independent cloning methods^3, 4^ and *in vitro*^*5*^*-* or *in vivo*^*6*^*-*homology-based methods, it is possible to construct almost any synthetic gene, biosynthetic pathway or genome. Of the many methods, the one-pot restriction-ligation method, Golden Gate Assembly^7^ (GGA), has become a standard method for the synthetic biology workflows due to its utility for rapid and high-throughput cloning. GGA utilizes type IIS restriction enzymes that cut distal to their recognition site, which allows overhangs to be designed with arbitrary sequence for scarless cloning. Although DNA synthesis technologies have advanced drastically, the synthesis of DNA sequences with high or low GC-content, stable secondary structures and multiple repeats remain challenging^8^, where especially repetitive DNA sequences result in synthesis failure or delays by vendors^9^. To circumvent some of these challenges, it is possible to codon optimize or codon scramble the sequence of genes encoding proteins to enable successful synthesis with standard methods^10^. However, this approach is not applicable for genes encoding repeats of functional non-coding RNA elements, since even a few sequence changes may change the folding and negatively affect the function of the RNA element. Utilization of functional RNA elements, either isolated from nature or *de novo* designed, has been increasingly popular in both the fields of synthetic biology^11^ and RNA nanotechnology^12^. Thus, a DNA assembly method to construct repetitive sequences is needed for successful gene construction.

Methods to assemble synthetic DNA with repeats have been developed. These techniques assemble shorter DNA sequences into larger repetitive DNA sequences using either restriction-ligation with type IIS restriction enzymes in multiple steps^13, 14^ or simply by annealing oligos^15^ to create sequence-defined overhangs for ligation. While these methods have proven useful for the scarless construction of repetitive DNA, the approach with type IIS restriction enzymes can be put into a GGA workflow for rapid assembly with minimal handling. GGA-based methods exist for the construction of at least 12 repeats for CRISPR guide-RNA arrays^16^. However, these methods rely on pre-constructed templates for the amplification of the scaffold part of the guide-RNA and are therefore not generalizable. Furthermore, these methods have not demonstrated the assembly of repetitive DNA, where specific linker or scaffold sequences with variable lengths are interspersed between the repeats such as for genes encoding RNA origami nanostructures^12^ that display multiple copies of the same RNA element (Figure 1). Therefore, we wanted to build on this prior work and make a DNA assembly method to solve this challenge.

**Figure 1.**
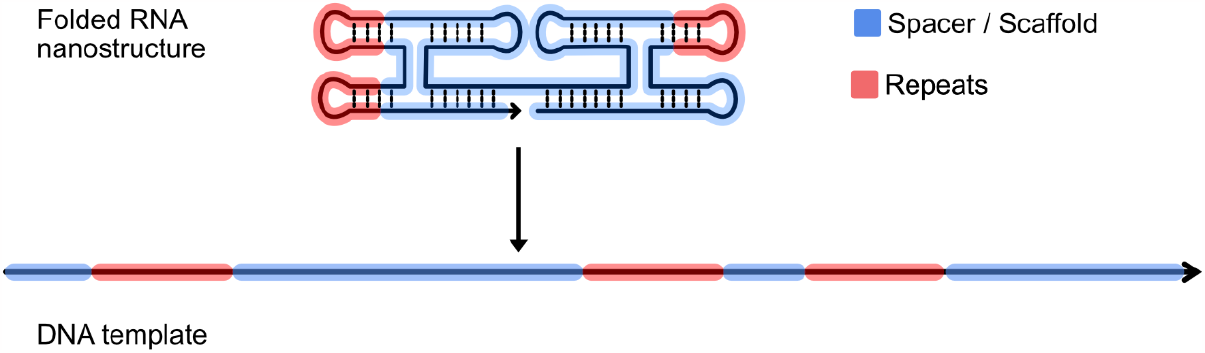
Example of repetitive non-coding RNA gene. Schematic representation of the secondary structure of a 2-helix RNA origami scaffold^12^ (blue) with repeated elements (red) and its associated DNA template.

## Results and discussion

Here, we describe a two-step DNA assembly method for the construction of synthetic genes with repetitive elements that vendors could not provide. An overview of the method can be seen in Figure 2. The method consists of splitting the gene of interest *in silico* into synthons of up to 80 base pairs (bp) using Molecular Biology design software such as Benchling^17^. The genes are split into roughly equal-sized fragments depending on the sequence size starting from position 1 to position 70-80 and then from position 70-80 to position 140-150 and so on. We chose to design our method around GGA since this method uses small overhangs (4 bp) that can be designed to minimize undesired annealing in contrast to homology-based approaches which require at least 15-bp homology^4^ and the synthons can be directly inserted into a desired destination vector. We used the GGA tool from Benchling to design compatible 4-bp overhangs. Other tools for overhang design exist such as NEB Golden Gate tool^18^ or the Golden Gate overhang designer from the Edinburgh biofoundry^19^. Synthons were made by oligo extension with a DNA polymerase using oligos of up to 60 nucleotides (nt) that overlap with 18-25 bases in their 3’-ends for hybridization to reduce misannealing and allow for extension of short oligos to longer synthons. The synthetic gene is finally assembled and inserted into a desired entry vector by GGA.

**Figure 2.**
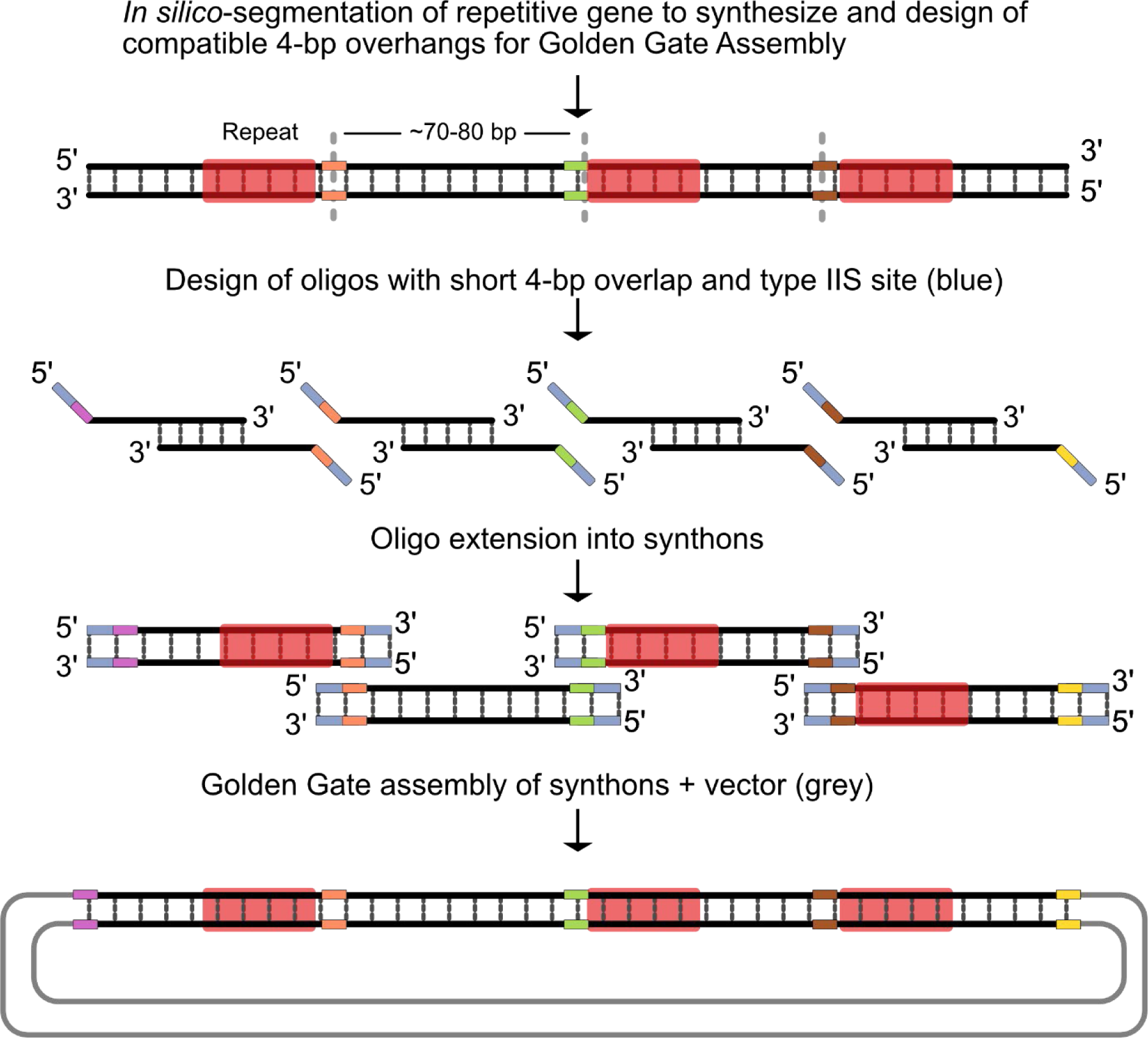
Overview of the two-step enzymatic synthesis of a synthetic gene. The designed gene with repeats (red) is split into 70-80 bp synthons *in silico* with 4-bp regions (orange, green, brown) that are used as overhangs for GGA. Oligos up to 60 nts are designed to cover the gene fragment and the oligos contain type IIS restriction enzyme sites in the 3’-ends (blue). Oligos covering the termini of the genes include overhangs (purple and yellow) compatible with an entry vector. Double-stranded DNA synthons are synthesized by oligo extension and inserted into a vector by GGA.

To demonstrate the applicability of our method, we assembled synthetic genes encoding tandem repeats of RNA aptamers with unique linker sequences between each repeat and RNA nanostructures with multiple repeated aptamers. These RNA constructs displayed different combinations of the known fluorogenic aptamers (Broccoli^20^, Corn^21^, Pepper^22^), the protein-binding aptamers (MS2^23^ hairpin and Cas6f^24^ hairpin) and a recently selected aptamer (P6Ba) that targets the SARS-CoV2 spike protein^25^. Details regarding their secondary structures (Figure S1) and sequences can be found in the supplementary material.

We chose these synthetic constructs because they require a well-defined sequence in between the repetitive elements and therefore cannot be synthesized by methods used for concatemerization of repeats^14^. Sequences ranging from 133 bp to 456 bp were designed. The sequences all failed initial screening for synthesis by the vendors Twist Bioscience and Integrated DNA Technologies^26^ (IDT) due to their content of repeated elements with repeat lengths ranging from 20 bp to 81 bp. Furthermore, all the sequences have a complexity score from IDT ranging from 30 to 139, well above 10 (Table 1), which is the maximum score allowed for synthesis. The sequence with the size of 133 bp was split into two fragments, the sequences ranging from 256 bp to 312 bp were split into four fragments, the sequence with 399 bp was split into five fragments and the two sequences with the sizes of 452 bp and 456 bp were split into six fragments. To synthesize the constructs, we used oligos of up to 60 nt, as this is the lowest price tier from IDT and each synthon was synthesized in individual reactions to reduce misannealing of the oligos that can cause faulty assembly. The synthons were assembled by GGA into different destination vectors. The constructs 2H-3xCorn and 2H-3xBroccoli were assembled with a linearized backbone, while the remaining constructs were assembled with a circular backbone. This was done due to the lack of availability of a desired circular entry backbone for those specific constructs. Plasmid sequences with accompanying sequencing data can be found in Table S2 and the sequences for the genes, synthons and oligos can be found in Table S3, Table S4, and Table S5, respectively.

**Table 1.** Summary of assembly fidelity for our eight different constructs with complexity scores from IDT.

| Name | Length (bp) | Synthons | Complexity score | Fidelity (correct out of total) |
| --- | --- | --- | --- | --- |
| 2x(Cas6f-hp-MS2wt) | 133 | 2 | 30.2 | 87.50% (7 out of 8) |
| 2H-3xCorn | 256 | 4 | 51.4 | 20.00% (2 out of 10) |
| 8xCas6f-hp | 280 | 4 | 139.2 | 62.50% (5 out of 8) |
| 2H-3xBroccoli | 301 | 4 | 103.3 | 62.50% (5 out of 8) |
| 8xMS2 | 312 | 4 | 101 | 87.50% (7 out of 8) |
| PX-Tri-P6Ba-UU | 399 | 5 | 88 | 25.00% (2 out of 8) |
| dAF14 | 452 | 6 | 123.9 | 12.50% (1 out of 8) |
| dAF10 | 456 | 6 | 112.9 | 12.50% (1 out of 8) |

We successfully constructed all the synthetic genes that could not be synthesized by the vendors (Table 1). For each construct, we sequenced at least eight clones with fidelity reported as the percentage of correct clones (Table 1). The highest observed fidelity of 87.5 % i.e., 7 correct clones out of 8 sequenced was observed for the assembly of 2x(Cas6f-hp-MS2wt) and 8xMS2.

We observed that the synthesis fidelity decreased with increasing number of synthons. The 4-synthon constructs were assembled with a fidelity higher than 60 % except for one construct, 2H-3xCorn, where only 20 % of the selected clones were successful after two separate cloning attempts. For 2H-3xCorn, it appears that the linear backbone self-ligated, which resulted in colonies that only propagated the vector (Table S1). We observed that failed assemblies had a lack of partial or whole synthons (Table S1). The lack of partial synthons indicates errors in the oligo extension that results in shorter synthons while the lack of whole synthons indicates that the overhangs are suboptimal for GGA. This could potentially be improved with computational tools for data-optimized design of Golden Gate overhangs^27^. The failed assemblies could also be due to misannealing of the wrong synthons during the GGA reaction or due to manual mishandling of synthons when preparing the assembly in the lab e.g., using the wrong concentrations. We also observed failed assemblies with just single-bp mutations or deletions. These mutations or deletions could be either due to misannealing of oligos during the extension step, or due to synthesis errors. Prior work has shown that assembly of synthetic genes with chemically synthesized oligos can be improved after gel-purifying the oligos^28^. Furthermore, we did not sequence verify any synthons before assembling the final synthetic genes. Therefore, errors in the synthons could also have occurred during the oligo extension. However, by relying only on cheap and relatively short synthetic DNA oligos that have a short turnaround time, it is possible to synthesize genes faster compared to synthetic gene synthesis and delivery from commercial vendors.

Taken together, using simple design tools we demonstrate the construction of synthetic genes of up to 456 bp with repetitive elements that are not possible for common vendors to manufacture through their standard platform due to the sequence complexity. We report that the assembly fidelity dropped below 50 % with an increase in synthon number and complexity, low fidelity should not be a concern in cases where gene synthesis is otherwise impossible and efficiency levels should still be financially viable and non-laborious even at higher synthon numbers.

While this method was developed to facilitate our own specific research needs in the construction of genes encoding synthetic non-coding RNA devices and RNA nanostructures with repetitive elements, we envision it to not only be of use for RNA synthetic biology and RNA nanotechnology, but ultimately an important addition to the molecular cloning toolbox.

## Materials and Methods

### RNA construct design

2x(Cas6f-hp-MS2wt), 8xCas6f-hp, 8xMS2 were all designed using the NUPACK design package. The secondary structures together with the sequences for the hairpins were used as design constraints. Linker sequences were designed as non-repetitive sequences. The design with the lowest ensemble score was chosen for synthesis. 2H-3xCorn, 2H-3xBr, PX-Tri-P6Ba-UU, dAF14, and dAF10 were all designed using the ROAD^12^ software and method for RNA origami design.

### Design of fragments and oligos

Genes of interest are divided into 70-80 bp fragments *in silico* using Benchling. Compatible 4-bp overhangs and initial primers with restriction site are designed with the Golden Gate assembly tool from Benchling. Initial designed oligos are then manually extended in the Benchling sequence editor to overlap by 18-25 bp for optimal annealing. Oligos are kept at a maximum length of 60 bases.

### Oligo extension for dsDNA synthesis

DNA oligos (Integrated DNA technologies) for synthons were ordered resuspended in 1x IDT TE buffer. Synthons were assembled by oligo extension using Q5 DNA Polymerase (NEB) in 25 µL reactions consisting of 1x Q5 reaction buffer, 200 µM of each dNTP, 0.5 unit Q5 DNA polymerase, 200 nM oligos (100 nM of each). Thermocycling was performed as following: initial denaturation at 98 °C for 30 s, then six cycles of: denaturation at 98 °C for 10 s, annealing at synthon-dependent temperatures for 20 s, elongation at 72 °C for 10 s, ending with a final elongation at 72 °C for 2 minutes. Synthons were purified with the Nucleospin Gel and PCR clean-up kit (Macherey-Nagel) following the manufacturer’s protocol using a vacuum manifold. Synthon concentration was measured with a Denovix-11. Purified synthons were diluted to 100 nM.

### Golden Gate reaction

Golden Gate reactions were performed with equimolar amounts of DNA using 25 or 50 femtomoles of synthon DNA, 25 femtomoles vector DNA, 0.5 µL T4 DNA ligase (NEB), 0.5 µL of either Esp3I (NEB) or BsaI (NEB) in 1x T4 DNA ligase buffer with 10 µM ATP (NEB) in 10 µL reactions. Golden Gate reactions consisted of 10 min at 37 °C, followed by 15 cycles of 5 min at 37 °C and 5 min at 16 °C followed by heat-inactivation of the enzymes by a 5 minutes incubation at 50 °C and at 80 °C.

The whole reaction was transformed into NEB Turbo cells following standard protocols and cells were plated on LB-agar plates containing either 100 µg/mL carbenicillin or 34 µg/ml chloramphenicol. Eight to ten non-fluorescent colonies for each construct were picked for plasmid propagation. Plasmids were propagated in cultures of terrific broth medium with appropriate antibiotics overnight at 37 °C and purified with NucleoSpin miniprep kit (Macherey-Nagel) following the manufacturer’s prototol. Plasmids were verified by Sanger sequencing (Eurofins Genomics).

## Acknowledgments

This project was financed by a Novo Nordisk Foundation Ascending Investigator grant (0060694). We thank Cody Geary for designing the PX-Tri-6Ba construct for our test. Thanks to Rita Rosendahl and Claus Bus for technical assistance.

