## Supplemental Material for "Enzymatic assembly of small synthetic genes with repetitive elements"

In addition to the material presented we also provide a link to a Benchling folder with all sequences and related information here - [Nguyen et al supplementary material (Benchling)](https://benchling.com/s/seq-KZtsrmvnzZTAjv6hVJnM)

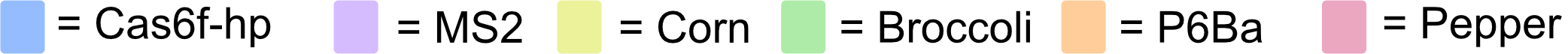

**2x(Cas6f-MS2)**

CCCGAAACUACUGCCGUAUAGGCAGACGCCCUACUAACUAACAUGAGGAUUACCCAUGUGCCCAAUCUCUACCACUGCCGUAUAGGCAGACUCCACUAACAUCGACAUGAGGAUUACCCAUGUACCCAGCCCC

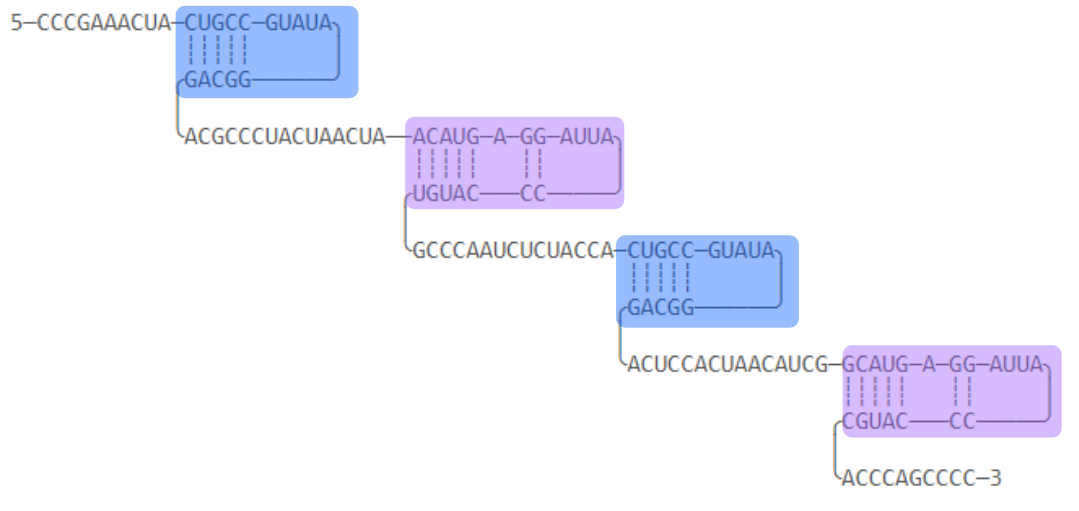

**2H-3xCorn**

GGACUCCAAUGGCCUGGCGCCUUCGGGUGUCAGGCGCGUUUGGCCGAGGAAGGAGGUCUGAGGAGGUCACUGGUCAAACGCCUAUGUGAAUUCGACACACAUAGCAUUGGGGUCCGUGUACGGCUUGUGGUGAAGUCGAAACACCACGUGUAGAGGCGAGGAAGGAGGUCUGAGGAGGUCACUGCCUUUAUACCUAACGCAACGAGGAAGGAGGUCUGAGGAGGUCACUGUUGCGUUGGAAGCCGUGCACCUGCCA

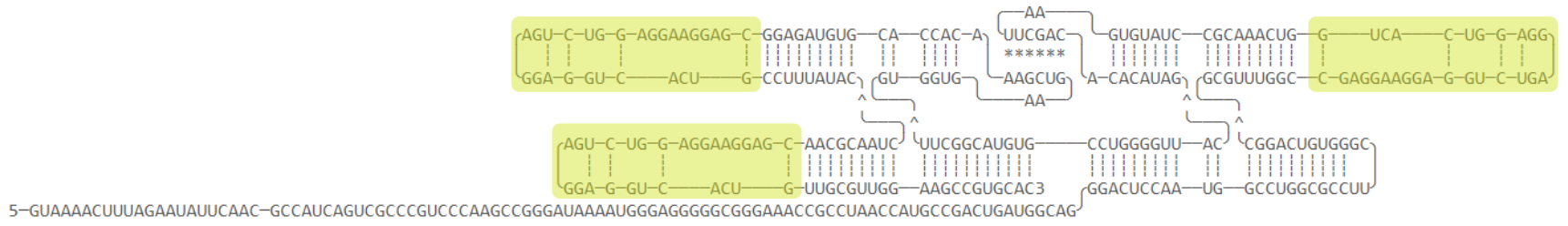

Figure S1. Secondary structure diagrams and sequences of the RNA structures. (*) represents kissing loop base pairs. Aptamers er colored. Figure continues on the following pages.

**8xCas6f**

CUGCCGUAUAGGCAGACACCUCCACUACCCUCACCCUGCCGUAUAGGCAGACCACUCCACCUUACACCUACUGCCGUAUAGGCAGACCACUCCAUACCACUCCAACUGCCGUAUAGGCAGCCUCACAUCCUCACCUACCCCUGCCGUAUAGGCAGACACCUCCACUCCACUCCCACUGCCGUAUAGGCAGACUCACUACCUCCACUCCCACUGCCGUAUAGGCAGACUCACUACCUCCACUCCCACUGCCGUAUAGGCAGACUCACUACCUCCACUCCCA

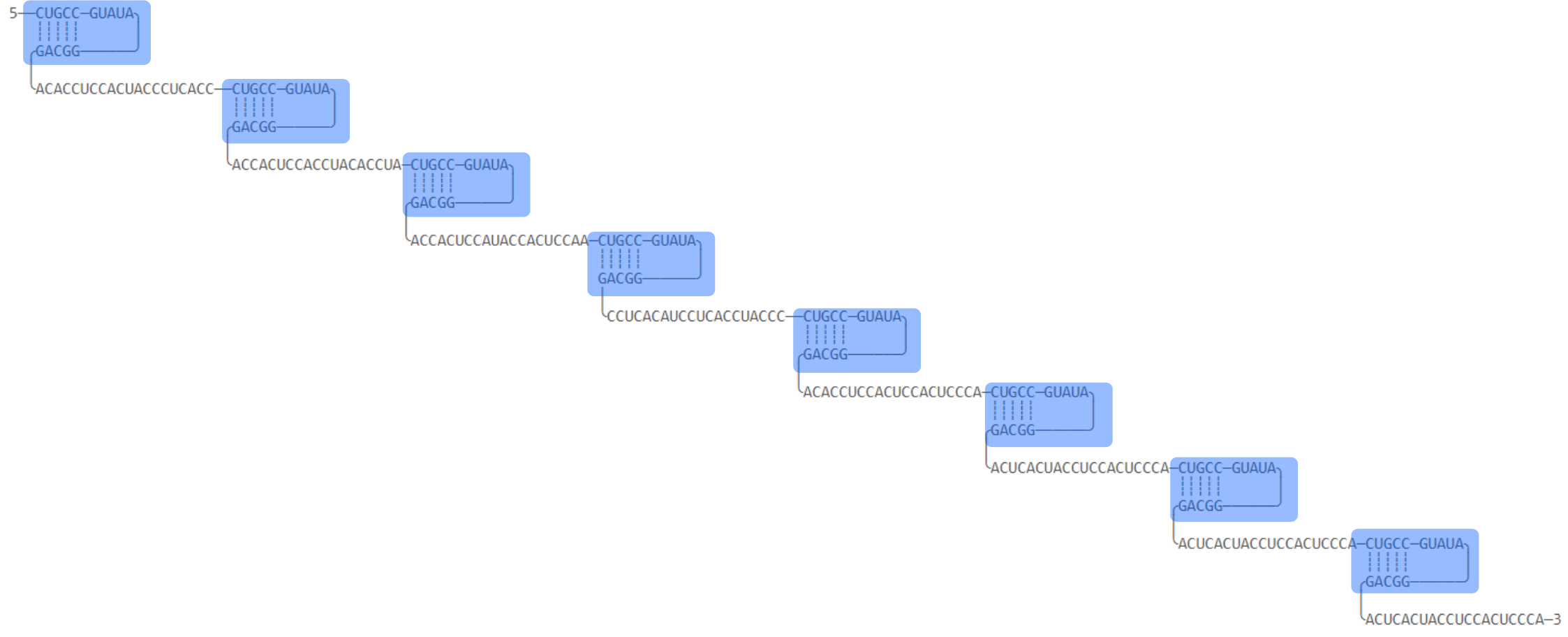

**2H-3xBroccoli**

GGACGUCGGGGAGGGUUAUCUUUCGAGAUGACCCUGGCACAGGGAGACGGUCGGGUCCAGUUCGCUGUUGAGUAGUGUGUGGGCUCCCUGUGUCUCAGAGAAAGACCGCAUCUCUGACCUCGACGUCCGCGGGAGGGAUCGGCGCAAGCGGUCAGCGCCGGUAAGGUGGAGACGGUCGGGUCCAGGGGUUCGCCCCUGUCGAGUAGAGUGUGGGCUCCAUCUUACGACGUUAGGAGACGGUCGGGUCCAGUUCGCUGUUGAGUAGUGUGUGGGCUCCUAGCGUCAUCUCUCCCGCCUGCCA

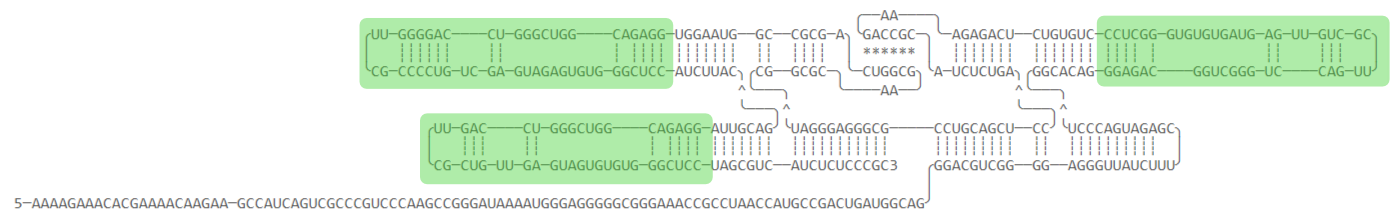

**Figure S1. (Continued)**

**8xMS2**

GCAUGAGGAUUACCCAUGCUACUUUAACUCCACUCCACCGUAUGAGGAUUACCCAUACAACUACCUACCUACCACUCCCGAUGAGGAUUACCCAUCGACUCUACCUCCACUACUUCCGCAUGAGGAUUACCCAUGCAUCCACUCCCACUCCAACUCGCAUGAGGAUUACCCAUGCACCCUACCUACCAAUCACCAUCAUGAGGAUUACCCAUGAACGCCUAUAACUACCACUCCCGAUGAGGAUUACCCAUCGAUACCACUACUCACUCCCAAGCAUGAGGAUUACCCAUGCACUCACCCUACAUCACCCUC

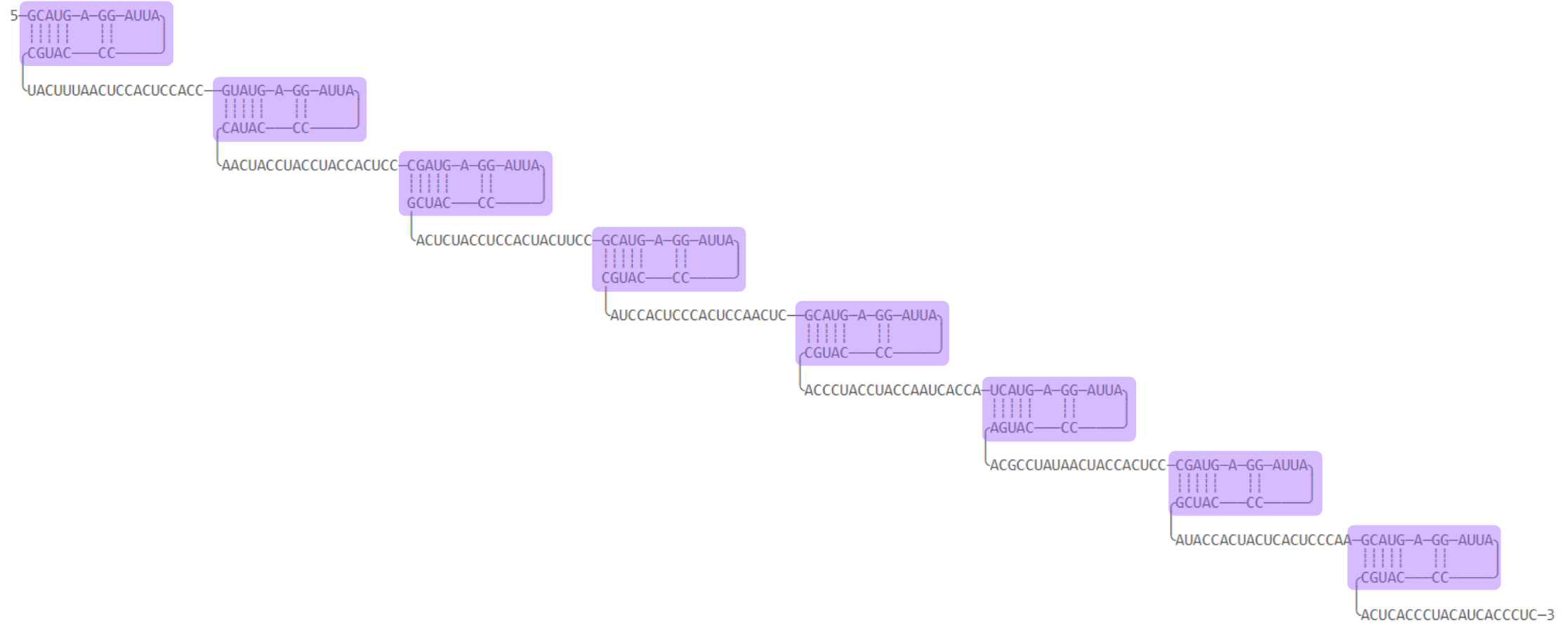

**PX-Tri-P6Ba-UU**

GGAAGGUGGUUGGCGACAUUUGUAAUUCCUGGACCGAUACUUCCGUCAGGACAGAGGUUGCCUUGGCAAGAUCCCUCAUUGGCGACAUUUGUAAUUCCUGGACCGAUACUUCCGUCAGGACAGAGGUUGCCUUCCAUCCAUCUUGCCUUGGCGACAUUUGUAAUUCCUGGACCGAUACUUCCGUCAGGACAGAGGUUGCCUUCCACCUUCCACGACACGGAAAGGGCUAUUCGUAGUCCUUUCCCUUCGUCAGGAUGGAUCUACCGGUGUUCGCAUCGGUAGAUUGAGGGUGACGAAGAAUAUCGGAGGUGUUCGCGCCUUCGAUGUUGUGUCGU

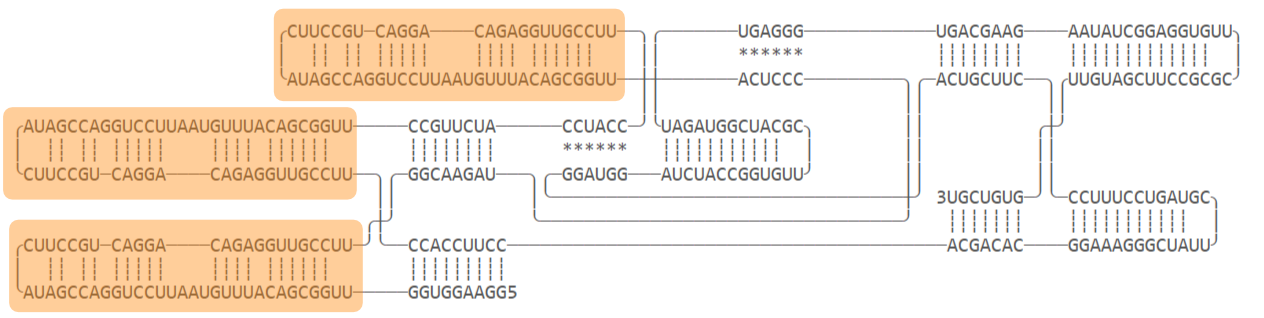

**Figure S1. (Continued)**

**dAF14**

GGAUACGUCUACGCUCAGUGCGUGUUGGUUCCCCAAUCGUGGCGUGUCGGCCUGCUUCGGCAGGCACUGGCGCCGGGAGCCAACGCUGCGUCGAAUCGCGCACGACGCACCACACUCGAAAGGGCAACGAGUGUCCGCUUUCGGCGUAAGAAGGGUGGGCCUCCAAUGCCCUAGGAGGCGGCAGGAAGCGCGAACCUGCCCUGGGUGCGCAGACGGUCGGGUCCAGUUCGCUGUCGAGUAGAGUGUGGGCUGCGCACCUAGCCUACUAGAUUCGUCUAGUAUGUGACGAGCAGACGGUCGGGUCCAGUUCGCUGUCGAGUAGAGUGUGGGCUGCUUGUCACAACCCUUCUUGCGUCGAAAGCGGGAGCGUCAUCCCCAAUCGUGGCGUGUCGGCCUGCUUCGGCAGGCACUGGCGCCGGGAUGGCGCUCGGGCACUGGGUGUGGACGUAUCC

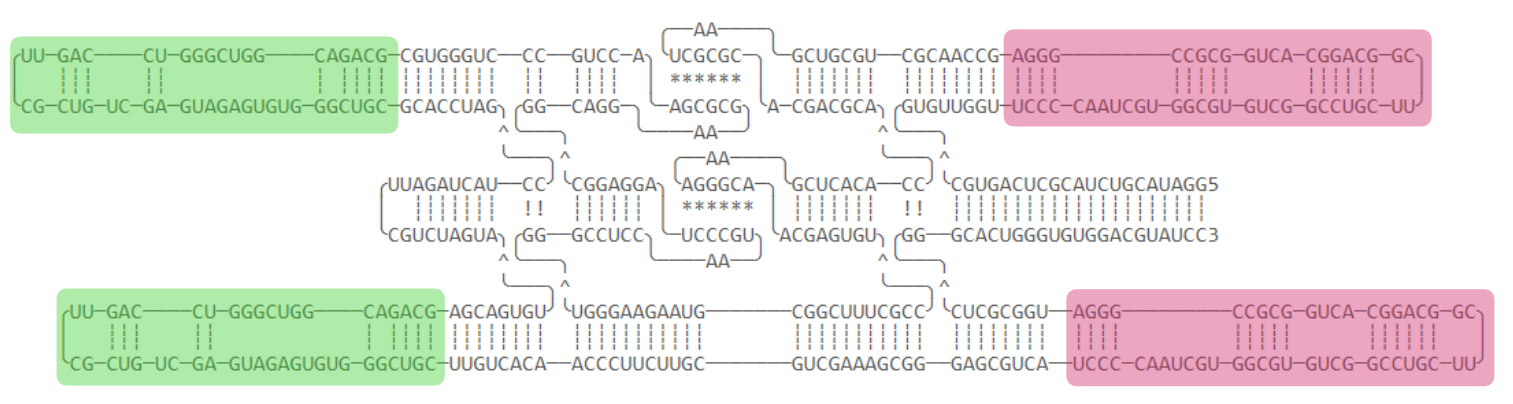

**dAF10**

GGAUACGUCUACGCUCAGUGAGCCGGAGUUGUCCCCAAUCGUGGCGUGUCGGCCUGCUUCGGCAGGCACUGGCGCCGGGACGACUCCGCUUCUGGCGCAGACGGUCGGGUCCAGUUCGCUGUCGAGUAGAGUGUGGGCUGCGCUAGAAGGGCUCCGAACUGAUCACGGAGCCGCGUCGCCUAAACCUACAAGGCGACUGGGAGUUUCUGGGACGUGGGACGGGAUCCAAGUAGGUAGGAUCCGUGCCUAAGAUCAGAAGGCACGACUGGUUGCAGACGGUCGGGUCCAGUUCGCUGUCGAGUAGAGUGUGGGCUGCAACUAGUCCGCUCGUUGCUCCCCAAUCGUGGCGUGUCGGCCUGCUUCGGCAGGCACUGGCGCCGGGAGCAGCGAGAUAGAUAACUUCGGUUAUCUAUUCCUACGUCCUAGAAACUUCCAUCACUGGGUGUGGACGUAUCC

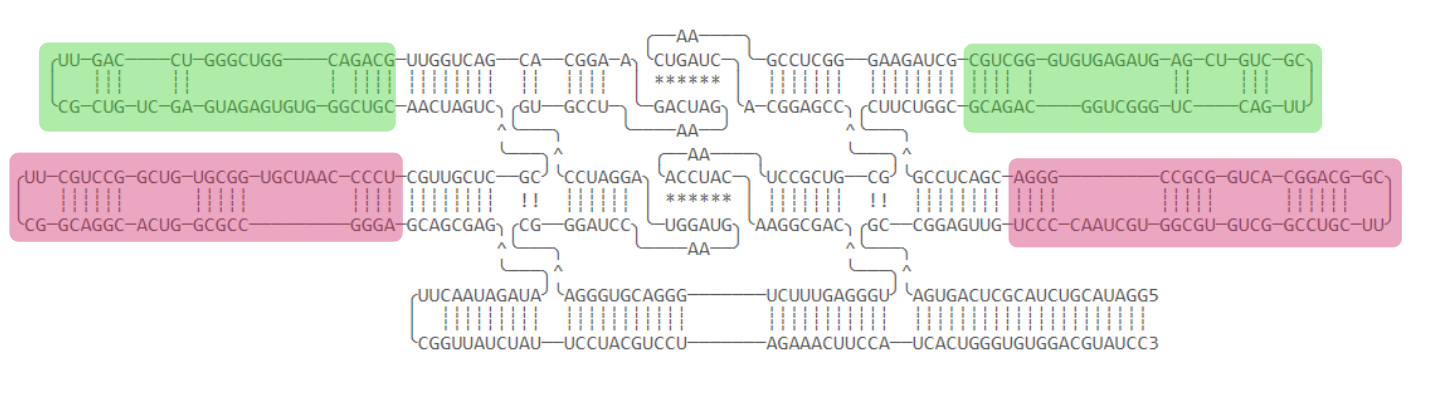

**Figure S1. (Continued)**

### Table S1: Summary on assembly fidelity with the comments on assembly failures.

| **Gene name** | **Length (bp)** | **Synthons** | **Fidelity** | **Comments** |
| --- | --- | --- | --- | --- |
| 2x(Cas6f-hp-MS2wt) | 133 | 2 | 87.50% | Several mutations in synthon 2 |
| 2H-3xCorn | 256 | 4 | 20.00% | All failures was vector ligation of the linearized backbone |
| 8xCas6f-hp | 280 | 4 | 62.50% | Duplication of synthon 3, several errors in synthon 4 |
| 2H-3xBroccoli | 301 | 4 | 81.82% | Lack of half synthon 1 and all of synthon 2-3. One failed assemble due to single bp deletion. One failed due to 1-bp and 2-bp deletions in synthons. |
| 8xMS2 | 312 | 4 | 87.50% | 1-bp mutation in synthon 2 |
| PX-Tri-P6Ba-UU | 399 | 5 | 25.00% | Lack of one to three synthons in complete sequence |
| dAF14 | 452 | 6 | 12.50% | Five out of eight failed assemblies were missing synthons. The remaining two had single bp deletions. |
| dAF10 | 456 | 6 | 12.50% | Missing a few synthons in the failed assemblies |

### Table S2: List of fully assembled DNA constructs and links to their plasmid sequence.

| **Gene name** | **Benchling link** |
| --- | --- |
| 2x(Cas6f-hp-MS2wt) | [Link to 2x(Cas6f-hp-MS2wt)](https://benchling.com/s/seq-6dLWJTezpcJyF5vlchfe?m=slm-lo9Cx1Zll7IRnrQaiwVn) |
| 2H-3xCorn | [Link to 2H-3xCorn](https://benchling.com/s/seq-nw4rZJo9SHzJQXSj8Sjx?m=slm-aE3yNYWV4VGmzCaP4rBN) |
| 8xCas6f-hp | [Link to 8xCas6f-hp](https://benchling.com/s/seq-WCfddX9MfDao2HsM1RtM?m=slm-OzOCAHwpl38VQ6rC0vIi) |
| 2H-3xBroccoli | [Link to 2H-3xBroccoli](https://benchling.com/s/seq-wGe53jfHU5PTDCgHXqPD?m=slm-2oUCgRgPIYR4WIXEdd3u) |
| 8xMS2 | [Link to 8xMS2](https://benchling.com/s/seq-7uLv02bquz2iV02s9onu?m=slm-Eidhu5N0dM1ffpYgzn1f) |
| PX-Tri-P6Ba-UU | [Link to PX-Tri-P6Ba-UU](https://benchling.com/s/seq-QpKe5a3D3tZVcMZDHH0w?m=slm-GNnvu3let7JXBTFbsZiy) |
| dAF14 | [Link to dAF14](https://benchling.com/s/seq-FlHTqd96QRpiWiM9dRUL?m=slm-vtzdGlo8v63QIWRLFW51) |
| dAF10 | [Link to dAF10](https://benchling.com/s/seq-WFSn9n3pEilTqw9fssm6?m=slm-HFkcwbaVx8Ks5wxIMY4P) |

### Table S3: List of linear synthetic DNA gene and links to their sequence.

**2x(Cas6f-MS2) -** [**Benchling link to sequence**](https://benchling.com/s/seq-DBPaODK1rr3IjEXIb9Ir?m=slm-N1KnGKS0Lzk5fbGbRdDG)

CCCGAAACTACTGCCGTATAGGCAGACGCCCTACTAACTAACATGAGGATTACCCATGTGCCCAATCTCTACCACTGCCGTATAGGCAGACTCCACTAACATCGACATGAGGATTACCCATGTACCCAGCCCC

**2H-3xCorn –** [**Benchling link to sequence**](https://benchling.com/s/seq-HYOwdypGpH8tlyxEFrMI?m=slm-hHnbzGakpevsM0wJ7Q03)

GGACTCCAATGGCCTGGCGCCTTCGGGTGTCAGGCGCGTTTGGCCGAGGAAGGAGGTCTGAGGAGGTCACTGGTCAAACGCCTATGTGAATTCGACACACATAGCATTGGGGTCCGTGTACGGCTTGTGGTGAAGTCGAAACACCACGTGTAGAGGCGAGGAAGGAGGTCTGAGGAGGTCACTGCCTTTATACCTAACGCAACGAGGAAGGAGGTCTGAGGAGGTCACTGTTGCGTTGGAAGCCGTGCACCTGCCA

**8xCas6fhp –** [**Benchling link to sequence**](https://benchling.com/s/seq-YLXysDr6QjqxgLRpFQTE?m=slm-Z1FNZxwi41Oyu2nHMO32)

CTGCCGTATAGGCAGACACCTCCACTACCCTCACCCTGCCGTATAGGCAGACCACTCCACCTTACACCTACTGCCGTATAGGCAGACCACTCCATACCACTCCAACTGCCGTATAGGCAGCCTCACATCCTCACCTACCCCTGCCGTATAGGCAGACACCTCCACTCCACTCCCACTGCCGTATAGGCAGACTCACTACCTCCACTCCCACTGCCGTATAGGCAGACTCACTACCTCCACTCCCACTGCCGTATAGGCAGACTCACTACCTCCACTCCCA

**2H-3xBroccoli –** [**Benchling link to sequence**](https://benchling.com/s/seq-WFSn9n3pEilTqw9fssm6?m=slm-5kpsd9rm9LAerbTkRz3w)

GGACGTCGGGGAGGGTTATCTTTCGAGATGACCCTGGCACAGGGAGACGGTCGGGTCCAGTTCGCTGTTGAGTAGTGTGTGGGCTCCCTGTGTCTCAGAGAAAGACCGCATCTCTGACCTCGACGTCCGCGGGAGGGATCGGCGCAAGCGGTCAGCGCCGGTAAGGTGGAGACGGTCGGGTCCAGGGGTTCGCCCCTGTCGAGTAGAGTGTGGGCTCCATCTTACGACGTTAGGAGACGGTCGGGTCCAGTTCGCTGTTGAGTAGTGTGTGGGCTCCTAGCGTCATCTCTCCCGCCTGCCA

**8xMS2 –** [**Benchling link to sequence**](https://benchling.com/s/seq-KZtsrmvnzZTAjv6hVJnM?m=slm-Qs8ClN33onvhzFxzRevy)

GCATGAGGATTACCCATGCTACTTTAACTCCACTCCACCGTATGAGGATTACCCATACAACTACCTACCTACCACTCCCGATGAGGATTACCCATCGACTCTACCTCCACTACTTCCGCATGAGGATTACCCATGCATCCACTCCCACTCCAACTCGCATGAGGATTACCCATGCACCCTACCTACCAATCACCATCATGAGGATTACCCATGAACGCCTATAACTACCACTCCCGATGAGGATTACCCATCGATACCACTACTCACTCCCAAGCATGAGGATTACCCATGCACTCACCCTACATCACCCTC

**PX-Tri-P6Ba-UU –** [**Benchling link to sequence**](https://benchling.com/s/seq-Ll4uubxqKqCsQax9YcIn?m=slm-X15HSeNlxnz07kQB5vma)

TAATACGACTCACTATAGGAAGGTGGTTGGCGACATTTGTAATTCCTGGACCGATACTTCCGTCAGGACAGAGGTTGCCTTGGCAAGATCCCTCATTGGCGACATTTGTAATTCCTGGACCGATACTTCCGTCAGGACAGAGGTTGCCTTCCATCCATCTTGCCTTGGCGACATTTGTAATTCCTGGACCGATACTTCCGTCAGGACAGAGGTTGCCTTCCACCTTCCACGACACGGAAAGGGCTATTCGTAGTCCTTTCCCTTCGTCAGGATGGATCTACCGGTGTTCGCATCGGTAGATTGAGGGTGACGAAGAATATCGGAGGTGTTCGCGCCTTCGATGTTGTGTCGTATCCCGAGACCggatccactgg

**dAF14 –** [**Benchling link to sequence**](https://benchling.com/s/seq-z6Hhf2xJ06En4F6S5bRI?m=slm-C8tWzaQssUSHaHuX8Acz)

GGATACGTCTACGCTCAGTGCGTGTTGGTTCCCCAATCGTGGCGTGTCGGCCTGCTTCGGCAGGCACTGGCGCCGGGAGCCAACGCTGCGTCGAATCGCGCACGACGCACCACACTCGAAAGGGCAACGAGTGTCCGCTTTCGGCGTAAGAAGGGTGGGCCTCCAATGCCCTAGGAGGCGGCAGGAAGCGCGAACCTGCCCTGGGTGCGCAGACGGTCGGGTCCAGTTCGCTGTCGAGTAGAGTGTGGGCTGCGCACCTAGCCTACTAGATTCGTCTAGTATGTGACGAGCAGACGGTCGGGTCCAGTTCGCTGTCGAGTAGAGTGTGGGCTGCTTGTCACAACCCTTCTTGCGTCGAAAGCGGGAGCGTCATCCCCAATCGTGGCGTGTCGGCCTGCTTCGGCAGGCACTGGCGCCGGGATGGCGCTCGGGCACTGGGTGTGGACGTATCC

**dAF10 –** [**Benchling link to sequence**](https://benchling.com/s/seq-qTDbe2rhIWl1qErYPnaO?m=slm-5RWWX60n8hNJ4tVP0koR)

GGATACGTCTACGCTCAGTGAGCCGGAGTTGTCCCCAATCGTGGCGTGTCGGCCTGCTTCGGCAGGCACTGGCGCCGGGACGACTCCGCTTCTGGCGCAGACGGTCGGGTCCAGTTCGCTGTCGAGTAGAGTGTGGGCTGCGCTAGAAGGGCTCCGAACTGATCACGGAGCCGCGTCGCCTAAACCTACAAGGCGACTGGGAGTTTCTGGGACGTGGGACGGGATCCAAGTAGGTAGGATCCGTGCCTAAGATCAGAAGGCACGACTGGTTGCAGACGGTCGGGTCCAGTTCGCTGTCGAGTAGAGTGTGGGCTGCAACTAGTCCGCTCGTTGCTCCCCAATCGTGGCGTGTCGGCCTGCTTCGGCAGGCACTGGCGCCGGGAGCAGCGAGATAGATAACTTCGGTTATCTATTCCTACGTCCTAGAAACTTCCATCACTGGGTGTGGACGTATCC

### Table S4: List synthon sequences used to assemble each gene (5’ to 3’).

| **Synthon name** | **Synthon sequence** |
| --- | --- |
| 2x(Cas6f-hp-MS2wt)_S1 | CCGTCTCCatctCCCGAAACTACTGCCGTATAGGCAGACGCCCTACTAACTAACATGAGGATTACCCATGTGCCCAATCTCTACCACTGGAGACGG |
| 2x(Cas6f-hp-MS2wt)_S2 | CCGTCTCCCACTGCCGTATAGGCAGACTCCACTAACATCGACATGAGGATTACCCATGTACCCAGCCCCttagGGAGACGG |
| 2H-3xCorn_S1 | GCATGGTCTCGGGGACTCCAATGGCCTGGCGCCTTCGGGTGTCAGGCGCGTTTGGCCGAGGAAGGAGGTCTGAGGAGGCGAGACCATGC |
| 2H-3xCorn_S2 | GCATGGTCTCCGAGGTCACTGGTCAAACGCCTATGTGAATTCGACACACATAGCATTGGGGTCCGTGTACGGCTTGTGGTGGGAGACCATGC |
| 2H-3xCorn_S3 | GCATGGTCTCCGGTGAAGTCGAAACACCACGTGTAGAGGCGAGGAAGGAGGTCTGAGGAGGTCACTGCCTTTATACCTAAGGAGACCATGC |
| 2H-3xCorn_S4 | GCATGGTCTCCCTAACGCAACGAGGAAGGAGGTCTGAGGAGGTCACTGTTGCGTTGGAAGCCGTGCACCTGCCATGGAGACCATGC |
| 8xCas6fhp_S1 | GCGTCTCGatctCTGCCGTATAGGCAGACACCTCCACTACCCTCACCCTGCCGTATAGGCAGACCACTCCACCTTACACCTACTCGAGACGC |
| 8xCas6fhp_S2 | GCGTCTCGTACTGCCGTATAGGCAGACCACTCCATACCACTCCAACTGCCGTATAGGCAGCCTCACATCCTCACCTACCCCTCGAGACGC |
| 8xCas6fhp_S3 | GCGTCTCCCCCTGCCGTATAGGCAGACACCTCCACTCCACTCCCACTGCCGTATAGGCAGACTCACTACCTCCACTCCCACGGAGACGC |
| 8xCas6fhp_S4 | GCGTCTCCCCACTGCCGTATAGGCAGACTCACTACCTCCACTCCCACTGCCGTATAGGCAGACTCACTACCTCCACTCCCAttagGGAGACGC |
| 2H-3xBroccoli_S1 | GCATGGTCTCCCAGGGACGTCGGGGAGGGTTATCTTTCGAGATGACCCTGGCACAGGGAGACGGTCGGGTCCAGTTCGCTGTTGAGTAGTGGGAGACCATGC |
| 2H-3xBroccoli_S2 | GCATGGTCTCGAGTGTGTGGGCTCCCTGTGTCTCAGAGAAAGACCGCATCTCTGACCTCGACGTCCGCGGGAGGGATCGGCGCAAGCGGTCGAGACCATGC |
| 2H-3xBroccoli_S3 | GCATGGTCTCGCGGTCAGCGCCGGTAAGGTGGAGACGGTCGGGTCCAGGGGTTCGCCCCTGTCGAGTAGAGTGTGGGCTCCATCTTACGACGAGACCATGC |
| 2H-3xBroccoli_S4 | GCATGGTCTCCACGACGTTAGGAGACGGTCGGGTCCAGTTCGCTGTTGAGTAGTGTGTGGGCTCCTAGCGTCATCTCTCCCGCCTGCCATCGGAGACCATGC |
| 8xMS2_S1 | CCGTCTCGatctGCATGAGGATTACCCATGCTACTTTAACTCCACTCCACCGTATGAGGATTACCCATACAACTACCTACCTACCACTCCCGACGAGACGG |
| 8xMS2_S2 | CCGTCTCGCCGATGAGGATTACCCATCGACTCTACCTCCACTACTTCCGCATGAGGATTACCCATGCATCCACTCCCACTCCAACTCGCACGAGACGG |
| 8xMS2_S3 | CCGTCTCCCGCATGAGGATTACCCATGCACCCTACCTACCAATCACCATCATGAGGATTACCCATGAACGCCTATAACTACCACTCCCGGAGACGG |
| 8xMS2_S4 | CCGTCTCCTCCCGATGAGGATTACCCATCGATACCACTACTCACTCCCAAGCATGAGGATTACCCATGCACTCACCCTACATCACCCTCttagGGAGACGG |
| PX-Tri-P6Ba-UU_S1 | CCGTCTCCatctTAATACGACTCACTATAGGAAGGTGGTTGGCGACATTTGTAATTCCTGGACCGATACTTCCGTCAGGACAGGGAGACGG |
| PX-Tri-P6Ba-UU_S2 | CCGTCTCGACAGAGGTTGCCTTGGCAAGATCCCTCATTGGCGACATTTGTAATTCCTGGACCGATACTTCCGTCAGGACAGAGGCGAGACGG |
| PX-Tri-P6Ba-UU_S3 | CCGTCTCCGAGGTTGCCTTCCATCCATCTTGCCTTGGCGACATTTGTAATTCCTGGACCGATACTTCCGTCAGGACAGAGGTTGCCTTCCAGGAGACGG |
| PX-Tri-P6Ba-UU_S4 | CCGTCTCCTCCACCTTCCACGACACGGAAAGGGCTATTCGTAGTCCTTTCCCTTCGTCAGGATGGATCTACCGGTGTTCGCATCGGTAGATTGGGAGACGG |
| PX-Tri-P6Ba-UU_S5 | CCGTCTCCATTGAGGGTGACGAAGAATATCGGAGGTGTTCGCGCCTTCGATGTTGTGTCGTATCCCGAGACCggatccactggttagGGAGACGG |
| dAF14_S1 | CCGTCTCCatctGGATACGTCTACGCTCAGTGCGTGTTGGTTCCCCAATCGTGGCGTGTCGGCCTGCTTCGGCAGGCACTGGCGCCGGGAGGGAGACGG |
| dAF14_S2 | CCGTCTCCGGAGCCAACGCTGCGTCGAATCGCGCACGACGCACCACACTCGAAAGGGCAACGAGTGTCCGCTTTCGGCGTAAGAAGGGGAGACGG |
| dAF14_S3 | CCGTCTCCAAGGGTGGGCCTCCAATGCCCTAGGAGGCGGCAGGAAGCGCGAACCTGCCCTGGGTGCGCAGACGGTCGGGTCCAGTGGAGACGG |
| dAF14_S4 | CCGTCTCCCAGTTCGCTGTCGAGTAGAGTGTGGGCTGCGCACCTAGCCTACTAGATTCGTCTAGTATGTGACGAGCAGACGGTCGGGAGACGG |
| dAF14_S5 | CCGTCTCCGTCGGGTCCAGTTCGCTGTCGAGTAGAGTGTGGGCTGCTTGTCACAACCCTTCTTGCGTCGAAAGCGGGAGCGTCATCCCCAGGAGACGG |
| dAF14_S6 | CCGTCTCGCCCAATCGTGGCGTGTCGGCCTGCTTCGGCAGGCACTGGCGCCGGGATGGCGCTCGGGCACTGGGTGTGGACGTATCCttagCGAGACGG |
| dAF10_S1 | GCGTCTCGatctGGATACGTCTACGCTCAGTGAGCCGGAGTTGTCCCCAATCGTGGCGTGTCGGCCTGCTTCGGCAGGCACTGGCGAGACGC |
| dAF10_S2 | GCGTCTCGCTGGCGCCGGGACGACTCCGCTTCTGGCGCAGACGGTCGGGTCCAGTTCGCTGTCGAGTAGAGTGTGGGCTGCGCTAGAAGGGCTCGAGACGC |
| dAF10_S3 | GCGTCTCGGGCTCCGAACTGATCACGGAGCCGCGTCGCCTAAACCTACAAGGCGACTGGGAGTTTCTGGGACGTGGGACGGGATCCAAGTCGAGACGC |
| dAF10_S4 | GCGTCTCCAAGTAGGTAGGATCCGTGCCTAAGATCAGAAGGCACGACTGGTTGCAGACGGTCGGGTCCAGTTCGCTGTCGAGTAGAGTGTGGGAGACGC |
| dAF10_S5 | GCGTCTCGTGTGGGCTGCAACTAGTCCGCTCGTTGCTCCCCAATCGTGGCGTGTCGGCCTGCTTCGGCAGGCACTGGCGCCGGGAGCGAGACGC |
| dAF10_S6 | GCGTCTCGGGAGCAGCGAGATAGATAACTTCGGTTATCTATTCCTACGTCCTAGAAACTTCCATCACTGGGTGTGGACGTATCCttagCGAGACGC |

### Table S5: List of oligo sequences used for synthon synthesis.

| **Oligo name** | **Oligo sequence** |
| --- | --- |
| 2x(Cas6f-hp-MS2wt).FWD (1) | CCGTCTCCatctCCCGAAACTACTGCCGTATAGGCAGACGCCCTACTAACTAACATGAGG |
| 2x(Cas6f-hp-MS2wt).REV (1) | CCGTCTCCAGTGGTAGAGATTGGGCACATGGGTAATCCTCATGTTAGTTAGTAGGGCGTC |
| 2x(Cas6f-hp-MS2wt).FWD (2) | CCGTCTCCCACTGCCGTATAGGCAGACTCCACTAACATCGACATGAGGATTACC |
| 2x(Cas6f-hp-MS2wt).REV (2) | CCGTCTCCctaaGGGGCTGGGTACATGGGTAATCCTCATGTCGATGTTAGTG |
| 2H-3xCorn.FWD1 | GCATGGTCTCGGGGACTCCAATGGCCTGGCGCCTTCGGGTGTCAGGCGCGTTTGG |
| 2H-3xCorn.REV1 | GCATGGTCTCGCCTCCTCAGACCTCCTTCCTCGGCCAAACGCGCCTGACACC |
| 2H-3xCorn.FWD2 | GCATGGTCTCCGAGGTCACTGGTCAAACGCCTATGTGAATTCGACACACATAGC |
| 2H-3xCorn.REV2 | GCATGGTCTCCCACCACAAGCCGTACACGGACCCCAATGCTATGTGTGTCGAATTC |
| 2H-3xCorn.FWD3 | GCATGGTCTCCGGTGAAGTCGAAACACCACGTGTAGAGGCGAGGAAGGAGGTCTGAGG |
| 2H-3xCorn.REV3 | GCATGGTCTCCTTAGGTATAAAGGCAGTGACCTCCTCAGACCTCCTTCCTCG |
| 2H-3xCorn.FWD4 | GCATGGTCTCCCTAACGCAACGAGGAAGGAGGTCTGAGGAGGTCACTGTTGCGTTG |
| 2H-3xCorn.REV4 | GCATGGTCTCCATGGCAGGTGCACGGCTTCCAACGCAACAGTGACCTCC |
| 8xCas6f-hp.FWD (1) | GCGTCTCGatctCTGCCGTATAGGCAGACACCTCCACTACCCTCACCCTGCCGTATA |
| 8xCas6f-hp.REV (1) | GCGTCTCGAGTAGGTGTAAGGTGGAGTGGTCTGCCTATACGGCAGGGTGAGGGTA |
| 8xCas6f-hp.FWD.(2) | GCGTCTCGTACTGCCGTATAGGCAGACCACTCCATACCACTCCAACTGCCGTATAG |
| 8xCas6f-hp.REV (2) | GCGTCTCGAGGGGTAGGTGAGGATGTGAGGCTGCCTATACGGCAGTTGGAGTGGTATG |
| 8xCas6f-hp.FWD (3) | GCGTCTCCCCCTGCCGTATAGGCAGACACCTCCACTCCACTCCCACTGCCGTATAG |
| 8xCas6f-hp.REV (3) | GCGTCTCCGTGGGAGTGGAGGTAGTGAGTCTGCCTATACGGCAGTGGGAGTGGAG |
| 8xCas6f-hp.FWD (4) | GCGTCTCCCCACTGCCGTATAGGCAGACTCACTACCTCCACTCCCACTGCCGTATAG |
| 8xCas6f-hp.REV (4) | GCGTCTCCctaaTGGGAGTGGAGGTAGTGAGTCTGCCTATACGGCAGTGGGAGTGGAG |
| 2H-3xBroccoli.FWD1 | GCATGGTCTCCCAGGGACGTCGGGGAGGGTTATCTTTCGAGATGACCCTGGCACAGGGAG |
| 2H-3xBroccoli.REV1 | GCATGGTCTCCCACTACTCAACAGCGAACTGGACCCGACCGTCTCCCTGTGCCAGGGTC |
| 2H-3xBroccoli.FWD2 | GCATGGTCTCGAGTGTGTGGGCTCCCTGTGTCTCAGAGAAAGACCGCATCTCTGACCTCG |
| 2H-3xBroccoli.REV2 | GCATGGTCTCGACCGCTTGCGCCGATCCCTCCCGCGGACGTCGAGGTCAGAGATGCGGTC |
| 2H-3xBroccoli.FWD3 | GCATGGTCTCGCGGTCAGCGCCGGTAAGGTGGAGACGGTCGGGTCCAGGGGTTCGC |
| 2H-3xBroccoli.REV3 | GCATGGTCTCGTCGTAAGATGGAGCCCACACTCTACTCGACAGGGGCGAACCCCTGGACC |
| 2H-3xBroccoli.FWD4 | GCATGGTCTCCACGACGTTAGGAGACGGTCGGGTCCAGTTCGCTGTTGAGTAGTGTGTGG |
| 2H-3xBroccoli.REV4 | GCATGGTCTCCGATGGCAGGCGGGAGAGATGACGCTAGGAGCCCACACACTACTCAACAG |
| 8xMS2.FWD (1) | CCGTCTCGatctGCATGAGGATTACCCATGCTACTTTAACTCCACTCCACCGTATGAGGA |
| 8xMS2.REV (1) | CCGTCTCGTCGGGAGTGGTAGGTAGGTAGTTGTATGGGTAATCCTCATACGGTGGAGTGG |
| 8xMS2.FWD (2) | CCGTCTCGCCGATGAGGATTACCCATCGACTCTACCTCCACTACTTCCGCATGAGGATTA |
| 8xMS2.REV (2) | CCGTCTCGTGCGAGTTGGAGTGGGAGTGGATGCATGGGTAATCCTCATGCGGAAGTAGTG |
| 8xMS2.FWD (3) | CCGTCTCCCGCATGAGGATTACCCATGCACCCTACCTACCAATCACCATCATGAGGATT |
| 8xMS2.REV (3) | CCGTCTCCGGGAGTGGTAGTTATAGGCGTTCATGGGTAATCCTCATGATGGTGATTGGT |
| 8xMS2.FWD (4) | CCGTCTCCTCCCGATGAGGATTACCCATCGATACCACTACTCACTCCCAAGCATGAGG |
| 8xMS2.REV (4) | CCGTCTCCctaaGAGGGTGATGTAGGGTGAGTGCATGGGTAATCCTCATGCTTGGGAGTG |
| PX-Tri-P6Ba-UU FWD (1) | CCGTCTCCatctTAATACGACTCACTATAGGAAGGTGGTTGGCGACATTTGTAATTCCTG |
| PX-Tri-P6Ba-UU REV (1) | CCGTCTCCCTGTCCTGACGGAAGTATCGGTCCAGGAATTACAAATGTCGCCAAC |
| PX-Tri-P6Ba-UU FWD (2) | CCGTCTCGACAGAGGTTGCCTTGGCAAGATCCCTCATTGGCGACATTTGTAATTCCTG |
| PX-Tri-P6Ba-UU REV (2) | CCGTCTCGCCTCTGTCCTGACGGAAGTATCGGTCCAGGAATTACAAATGTCGCCAAT |
| PX-Tri-P6Ba-UU FWD (3) | CCGTCTCCGAGGTTGCCTTCCATCCATCTTGCCTTGGCGACATTTGTAATTCCTGGACCG |
| PX-Tri-P6Ba-UU REV (3) | CCGTCTCCTGGAAGGCAACCTCTGTCCTGACGGAAGTATCGGTCCAGGAATTACAAATGT |
| PX-Tri-P6Ba-UU FWD (4) | CCGTCTCCTCCACCTTCCACGACACGGAAAGGGCTATTCGTAGTCCTTTCCCTTCGTCAG |
| PX-Tri-P6Ba-UU REV (4) | CCGTCTCCCAATCTACCGATGCGAACACCGGTAGATCCATCCTGACGAAGGGAAAGGACT |
| PX-Tri-P6Ba-UU FWD (5) | CCGTCTCCATTGAGGGTGACGAAGAATATCGGAGGTGTTCGCGCCTTCGATGTTGTG |
| PX-Tri-P6Ba-UU REV (5) | CCGTCTCCctaaccagtggatccGGTCTCGGGATACGACACAACATCGAAGGCGCG |
| dAF14 FWD (1) | CCGTCTCCatctGGATACGTCTACGCTCAGTGCGTGTTGGTTCCCCAATCGTGGCGTG |
| dAF14 REV (1) | CCGTCTCCCTCCCGGCGCCAGTGCCTGCCGAAGCAGGCCGACACGCCACGATTGGGGAAC |
| dAF14 FWD (2) | CCGTCTCCGGAGCCAACGCTGCGTCGAATCGCGCACGACGCACCACACTCGAAAGG |
| dAF14 REV (2) | CCGTCTCCCCTTCTTACGCCGAAAGCGGACACTCGTTGCCCTTTCGAGTGTGGTGCGTCG |
| dAF14 FWD (3) | CCGTCTCCAAGGGTGGGCCTCCAATGCCCTAGGAGGCGGCAGGAAGCGCGAACCTG |
| dAF14 REV (3) | CCGTCTCCACTGGACCCGACCGTCTGCGCACCCAGGGCAGGTTCGCGCTTCCTGC |
| dAF14 FWD (4) | CCGTCTCCCAGTTCGCTGTCGAGTAGAGTGTGGGCTGCGCACCTAGCCTACTAGATTCG |
| dAF14 REV (4) | CCGTCTCCCGACCGTCTGCTCGTCACATACTAGACGAATCTAGTAGGCTAGGTGCGCAG |
| dAF14 FWD (5) | CCGTCTCCGTCGGGTCCAGTTCGCTGTCGAGTAGAGTGTGGGCTGCTTGTCACAACCCTT |
| dAF14 REV (5) | CCGTCTCCTGGGGATGACGCTCCCGCTTTCGACGCAAGAAGGGTTGTGACAAGCAGCCC |
| dAF14 FWD (6) | CCGTCTCGCCCAATCGTGGCGTGTCGGCCTGCTTCGGCAGGCACTGGCGCCGGGATG |
| dAF14 REV (6) | CCGTCTCGctaaGGATACGTCCACACCCAGTGCCCGAGCGCCATCCCGGCGCCAGTG |
| dAF10 FWD (1) | GCGTCTCGatctGGATACGTCTACGCTCAGTGAGCCGGAGTTGTCCCCAATCGTGGC |
| dAF10 REV (1) | GCGTCTCGCCAGTGCCTGCCGAAGCAGGCCGACACGCCACGATTGGGGACAACTCC |
| dAF10 FWD (2) | GCGTCTCGCTGGCGCCGGGACGACTCCGCTTCTGGCGCAGACGGTCGGGTCCAGTTCGC |
| dAF10 REV (2) | GCGTCTCGAGCCCTTCTAGCGCAGCCCACACTCTACTCGACAGCGAACTGGACCCGACCG |
| dAF10 FWD (3) | GCGTCTCGGGCTCCGAACTGATCACGGAGCCGCGTCGCCTAAACCTACAAGGCGACTGGG |
| dAF10 REV (3) | GCGTCTCGACTTGGATCCCGTCCCACGTCCCAGAAACTCCCAGTCGCCTTGTAGGTTTAG |
| dAF10 FWD (4) | GCGTCTCCAAGTAGGTAGGATCCGTGCCTAAGATCAGAAGGCACGACTGGTTGCAGACGG |
| dAF10 REV (4) | GCGTCTCCCACACTCTACTCGACAGCGAACTGGACCCGACCGTCTGCAACCAGTCGTGC |
| dAF10 FWD (5) | GCGTCTCGTGTGGGCTGCAACTAGTCCGCTCGTTGCTCCCCAATCGTGGCGTGTCG |
| dAF10 REV (5) | GCGTCTCGCTCCCGGCGCCAGTGCCTGCCGAAGCAGGCCGACACGCCACGATTGGG |
| dAF10 FWD (6) | GCGTCTCGGGAGCAGCGAGATAGATAACTTCGGTTATCTATTCCTACGTCCTAGAAACTT |
| dAF10 REV (6) | GCGTCTCGctaaGGATACGTCCACACCCAGTGATGGAAGTTTCTAGGACGTAGGAATAGA |
